# Preliminary evidence for upward elevational range shifts by Eastern Himalayan birds

**DOI:** 10.1101/2020.10.13.337121

**Authors:** Krishna S. Girish, Umesh Srinivasan

## Abstract

The ongoing climate crisis is one of the most significant threats to biodiversity globally. As the Earth warms, species have adapted by shifting their geographical ranges either polewards, or in mountainous regions, upslope towards higher elevations, presumably to continue to inhabit a suitable thermal environment. Upslope range shifts are of particular concern in tropical mountain ranges because: (a) tropical species are particularly thermally sensitive, (b) as species move upwards, they can run out of habitable space, leading to local extirpations, and (c) tropical mountains harbour a disproportionately high fraction of the planet’s terrestrial biodiversity – rapid upslope range shifts can therefore result in significant biodiversity losses. We used citizen science data over a 13-year period to evaluate whether 39 eastern Himalayan bird species might be shifting to higher elevations over time, by analyzing changes in the frequency of reporting of these species at birdwatching hotspots. For these species, we find evidence consistent with upslope range shifts, with species with the bulk of their elevational ranges below the hotspot elevation showing increases in their reporting frequency over time, and those with most of their elevational ranges above the hotspot elevation declining in reporting frequency. Our findings are suggestive of rapid responses to climate change by eastern Himalayan birds. We caution that eastern Himalayan bird species might be at special risk from increasing global temperatures because of their heightened thermal sensitivity coupled with particularly high rates of warming in the region. Eastern Himalayan birds are likely to require large tracts of undisturbed natural habitat across entire elevational gradients to be able to track changing temperatures by moving to higher elevations to remain resilient to climate change.

**SUMMARY:**

- One of the most fundamental responses of species to changing temperatures is to change their geographic ranges, possibly to track the range of temperatures that is ideal for their survival.
- With increasing climate warming, this shift could happen by moving towards the poles or in mountainous areas, towards the summit.
- Due to the high thermal sensitivity of tropical species and the decrease in space as species move up mountains, the extremely biodiverse bird communities of tropical mountains are particularly vulnerable.
- Using citizen science data from birding hotspots along an elevational gradient in the Eastern Himalayas over a 13-year period, we measured the change in reporting frequency of 39 common bird species.
- Changes in reporting frequency are generally consistent with the fact that upslope shifts are taking place in the Eastern Himalayas, similar to results for birds from other tropical mountains.

## INTRODUCTION

The sixth ever mass extinction in Earth’s history is currently underway (Ceballos & Ehrlich, 2018), and is driven primarily by the direct and indirect effects of climate change and the loss and degradation of natural habitats (Pimm 2008). Among the many impacts that climate change has on biodiversity, one of the most fundamental is the alteration of the geographical ranges of species (Intergovernmental Panel on Climate Change (IPCC), 2007; Chen et al., 2011). With climate change leading to ever-increasing temperatures across the globe, both terrestrial and marine species are shifting their ranges towards cooler regions, either latitudinally (i.e., towards the poles), or elevationally (i.e., towards mountain summits), or both (Parmesan & Yohe, 2003; Chen et al., 2009; La Sorte & Jetz, 2010, Lenoir et al., 2020). Upslope shifts in species’ elevational ranges, especially for thermally sensitive tropical species inhabiting relatively aseasonal habitats (Janzen, 1967; Wormworth & Şekercioğlu, 2011; Freeman et al., 2020), have been demonstrated for a range of biodiversity, from comparatively immobile groups such as plants (Feeley et. al., 2011; Morueta-Holme et. al., 2015; Salick et al., 2019) to more mobile taxa such as ectotherms (Raxworthy et al., 2008; Chen et al., 2009), birds (Şekercioğlu et al., 2008; Freeman & Freeman, 2014; Freeman et al., 2018), and mammals (Moritz et al., 2008; Rowe et al., 2015). Plants in the Himalayas have been shown to move upslope, presumably as a response to increasing temperature (Salick et al., 2019), and it is likely that Himalayan birds will track these vegetational and thermal shifts (Lenoir & Svenning, 2015), especially in the light of well-documented and rapid upslope shifts to higher elevations by montane birds in other tropical regions such as in Papua New Guinea and in the Peruvian Andes (Freeman & Freeman, 2014, Freeman et al., 2018).

These rapid upslope shifts present a problem for the continued survival of tropical montane species because the approximately conical structure of mountains causes area to decline with elevation, leaving progressively less habitat for species to colonise as they move summit-wards (but see Elsen & Tingley, 2015). This increases the probability of mountaintop local extinctions (Freeman et al., 2018), via not only a direct reduction in area available for occupancy, but also through indirect effects such as increased competition and other secondary and stochastic drivers of local extirpation, a phenomenon referred to as the “escalator to extinction” (Thomas et al., 2004; Marris, 2007; Brook et al., 2008; Şekercioğlu et al., 2012; Elsen & Tingley, 2015). Biodiversity losses arising from local extirpations on tropical mountains will be significant, because tropical mountains house a disproportionately high fraction of Earth’s terrestrial biodiversity and endemic species (Myers et al., 2000).

With rapidly accelerating warming, predicting how tropical montane species shift their ranges to higher elevations is essential: (a) because different species are likely to move upslope at different rates, leading to novel ecological communities and species interactions (Angert et al., 2011), and (b) to plan adequate long-term and large-scale measures to improve the resilience of species to climate change. Such an understanding is especially urgent for eastern Himalayan species, because the region is exceptionally biodiverse (second only to the Amazon-Andes system; Grenyer et al., 2006), and is warming three times as fast as the global average (Shrestha et al., 2012). Further, amongst various taxa, birds also provide a useful study system to investigate the impacts of climate change because they are relatively well-studied compared with other taxonomic groups, and citizen science initiatives have made large amounts of data collected at certain sites over the years publicly available. Finally, because the Himalayas are warming much faster than the global average, species’ responses to climate change in the Himalayas can serve as a bellwether to anticipate how tropical montane biodiversity might be impacted by climate change in the future.

Given the thermal sensitivity of tropical species in general (Janzen, 1967; Perez et al., 2016) and eastern Himalayan birds in particular (Srinivasan et al., 2018), and the rapidity of warming in the Eastern Himalayas (Sreshtha et al., 2012), we hypothesized that eastern Himalayan bird species would show rapid upslope elevational range shifts (see Srinivasan & Wilcove 2020). Using data from a relatively well-birded site in the eastern Himalayas, we compared the frequency with which species were recorded in eBird (eBird, 2020) checklists between two time periods (2006-2010 and 2016-1019) at two birding “hotspots” (at 1950 and 2450m above sea level (ASL); see Methods). We predicted that the difference in reporting frequency of species over time would be strongly related to whether species’ elevational ranges were largely above or below the elevation of these birdwatching hotspots. For species for which the mid-point of their elevational ranges were below the elevation of the hotspot, we expected increases in reporting frequency over time as species moved upslope and closer to the hotspot elevation. Species for which the mid-point of their elevational ranges were above the hotspot elevation should show movement upslope away from the hotspot, and therefore declines in reporting frequency at the hotspot over time.

## METHODS

### Study Areas

We studied the distribution of bird populations in Eaglenest Wildlife Sanctuary (27.07°N, 92.40°E; EWLS) and Singchung Bugun Village Community Reserve (27.15°N, 92.45°E; SBVCR), West Kameng district, Arunachal Pradesh, India. In the region as a whole, EWLS and SBVCR represent an area with minimal land-use change, apart from some historical military-related construction activity, which was halted several decades before, and low-intensity agriculture. EWLS and SBVCR also encompass vast, continuous elevational gradients, potentially allowing species to move upslope in response to climate change. It is therefore highly likely that any contemporary shifts in elevational ranges result from climate change.

We initially selected all five birdwatching hotspots designated by eBird within EWLS and SBVCR for analysis. eBird is an online real-time and freely downloadable citizen science database in which birdwatchers from across the world upload their sightings and records of birds. Some locations are particularly important for a suite of species, which are then designated hotspots. Within EWLS and SBVCR, the designated hotspots are Khellong (750m ASL), Sessni (1250m), Bongpu (1950m), Lama Camp (2350m) and Eaglenest Pass (2780m). Our aim was to compare how the proportion of checklists lists in which a species was recorded changed over time at a given hotspot (elevation). To do this, we had to:

a. Limit ourselves to analyzing data from only two hotspots (Bongpu and Lama Camp, montane broadleaved forest) that were more intensively birded and had enough checklists to compare reporting frequencies over time.
b. Limit ourselves to analyzing data from only stationary and complete checklists and *not* traveling checklists, because of the possibly significant elevational changes on traveling even short distances within the Himalayas.
c. Limit ourselves to data from the breeding season (March to May) to avoid the confounding impacts of seasonal movements (elevational migration) of birds within the Himalayas.

### Calculations: eBird Proportions

For each hotspot (Bongpu and Lama Camp), we split the available online checklists into two time periods spanning 14 years, based on the years from which the checklists were available – 2006 to 2010, and 2016 to 2019. No checklists satisfying our criteria were available for the years 2011 to 2015, or prior to 2006. For each species recorded at least ten times across checklists in each time period and in each hotspot, we calculated the proportion of checklists it was recorded in between 2006 and 2010 and then the proportion of checklists it was recorded in between 2016 and 2019, and subtracted the former from the latter. A positive number therefore means that the frequency of reporting of the species increased over time at the given elevation, and a negative number indicates that the frequency of reporting declined at the given elevation. We expected that for a species, the difference in the proportion of records over years would be correlated with the elevational range of a species relative to the elevation of the hotspot. Species with most of their historical elevational range below the hotspot elevation should show increases in reporting frequency as climate warming compels such species to move upslope to occupy higher elevations than before. Species with most of their elevational ranges above a given hotspot elevation should show declines in reporting frequency over time as they leave thermally suboptimal elevational zones for higher elevations. We extracted data on the midpoints of Himalayan species’ elevational ranges from the book Birds in Bhutan: Status and Distribution (Spierenburg, 2005), and for the species for which elevational data was not available from the book, we used the information given on the website Birds of the World (Billerman et al., 2020). We then used linear regression to correlate the changes in reporting frequency with the elevational distance of the midpoint of the species range from the hotspot. All analyses were done using Program R (R Core Team, 2020).

### Assumptions and Limitations of our Analyses

Our analyses assume several patterns, and are also limited by nature and quantity of the data we use for our inferences. We highlight some of these key cautions in this section, and hope that future research can overcome some of these limitations.

1. Abundant range centres: Our predictions relating to upward range shifts towards or away from elevational hotspots depending on the upper and lower elevational bounds of species’ range limits contain two related assumptions: (i) that elevational ranges are constrained by abiotic factors (mainly temperature) rather than biotic interactions such as competition (but see Freeman et al., 2019, and Burner et al., 2020), and will therefore respond primarily to changes in the thermal environment, and (ii) for any species, the relationship between elevation and abundance would show a unimodal distribution, such that abundance approaches zero at the upper and lower bounds of the elevational range of the species, and approaches a peak at some intermediate elevation (Gaston, 2009; Burner et al. 2019).
2. Data limitations:

i. Proportion data: The aim of our analyses was to infer elevational range shifts of Eastern himalayan bird species, likely in response to climate change. Truly measuring this would require showing significant changes in the lower and upper bound of species’ elevational ranges over time. These data, unfortunately, are not available, and we use elevation-specific changes in the proportion with which species are reported in checklists over time as a proxy for change in abundance with elevation, and therefore change in elevational range (see assumption #1 also). Further, we have no way of calculating the degree to which a given shift (in meters) in elevational range corresponds to a change over time in the proportion of checklists in which a species was recorded. Our response variable, therefore, is difficult to interpret biologically, and does not have clearly interpretable physical units.
ii. Volume of data: The number of checklists we use for our analyses are low – 59 for Bongpu and 115 for Lama Camp. A further limitation of our analyses arises from the fact that the checklists also have to be divided into two time periods, thereby reducing the number of checklists per time period. While the eastern Himalayas are exceptionally biodiverse, these low volumes of data have led to the unavoidable jettisoning of a majority of bird species from the analyses, limiting us to analyzing changes in reporting frequency for only 39 common species (out of 309 that were recorded in total in eBird checklists).
iii. Life-history traits: Because of the paucity of checklists and the low number of species included in our analyses, there is limited variation in life-history traits; we were therefore unable to relate variability in upward range shifts as a function of: (a) direct physiological mechanisms such as thermal tolerances by examining correlations between reporting frequency and thermally relevant species traits such as body mass, and/or (b) via indirect mechanisms such as the need to track food resources, by examining species’ diets and reporting frequencies (Angert et al., 2011).
3. Finally, we assumed that species did not become any more resilient to climate change and associated thermal stress over our study period.

## RESULTS

Across both hotspots, we extracted 3,966 individual records belonging to 309 species from eBird. Of these, most species were recorded infrequently, and had to be excluded from the analyses, leaving us with 1,821 individual records representing 39 relatively common species. These records were part of 59 checklists from Bongpu and 115 checklists from Lama Camp. The number of checklists from Khellong (04), Sessni (18) and Eaglenest Pass (21) were too low for analyses.

As expected, at both hotspots separately, species’ elevational ranges were correlated with changes in reporting frequency over time. At Bongpu (1950m, Figure 1A) and Lama Camp (2450m, Figure 1B), species with elevational ranges largely below the hotspot-specific elevations (i.e., positive values on the x-axis) showed increases in reporting frequency over time, while species that occurred largely above the elevation of the hotspot showed reporting declines (linear regression; *β*_Bongpu_ = 22% average change in reporting frequency per 1000m difference from mid-point of species’ elevational range, *F*= 4.51, df=1 and 27, *p* = 0.04, *R^2^* = 0.14; *β*_Lama Camp_ = 16% per 1000m, *p* = 0.02; *R^2^* = 0.15; *F* = 6.04, df=1 and 35). Some species had wide elevational ranges, and were found at both hotspots – for these species, patterns in reporting frequency differed between the two hotspots. For instance, the Chestnut-crowned Laughingthrush (*Trochalopteron erythrocephalum*), showed a marked ~40% decline in reporting frequency at 1950m (white dot in Figure 1A) but only a ~10% decline in reporting frequency at 2450m (white dot in Figure 1B).

**Figure 1.**
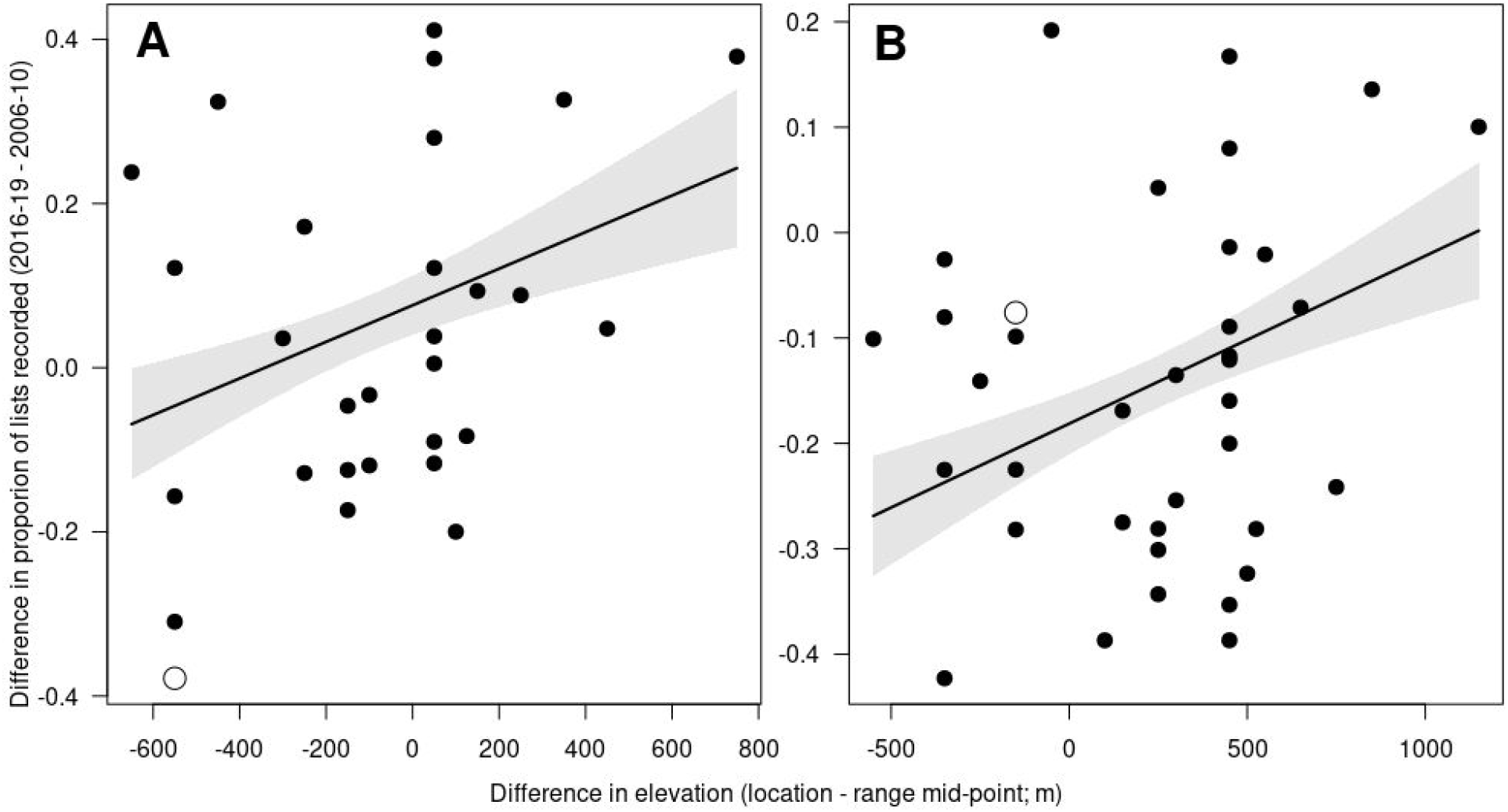
At both 1950m and 2450m ASL, species found at lower elevations (positive values on the x-axis) also had higher reporting frequencies in later eBird checklists (2016-19) than in earlier checklists (2006-2010). The open circle in both panels shows values for the Chestnut-crowned Laughingthrush (*Trochalopteron erythrocephalum*). At 2450m (B), which is roughly the mid-point of the elevational range for this species, there is a small decline (of ~10%) in reporting frequency over time. At 1950m (A), however, this species, with most of its elevational range above 1950m shows marked (~40%) decline in reporting frequency in eBird checklists over time, which is consistent with upslope range shifts.

## DISCUSSION

Our results from analyses of the change in the reporting frequencies of species from citizen science data are consistent with upslope range shifts in eastern Himalayan bird species. We find these patterns over a relatively short period (13 years) for a limited subset of species that we were able to include in the analyses, and with the use of reporting frequency data as a proxy for actual elevational range shifts, rather than measuring changes in the lower and/or upper bound of the elevational ranges of species over time. Our results relating species-specific elevational ranges and differences in reporting frequency over time are also consistent with recent long-term demographic work from the same study area (Srinivasan & Wilcove 2020) at one of the hotpots we analyze (Bongpu, 1950m ASL). At Bongpu, species with elevational ranges largely below 1950m showed survival increases over an eight-year period, while those with most of their elevational ranges above 1950m faced survival declines over time.

We caution that our results—although certainly expected in the light of rapid warming in the Eastern Himalayas (Sreshtha et al. 2012)—should be viewed as preliminary until such time that (a) direct measures of change in elevational range over time are available, (b) a larger number of species can be analyzed to examine variability in species’ responses to climate change, (c) more detailed analyses of long-term distribution patterns of mountain-top species can potentially identify local extirpations, and (d) across-Himalayan comparisons can be made to check for heterogeneity in species’ response to climate change along the length of the range. A large number of studies on the elevational ranges of Himalayan birds have already been carried out across the entire range (e.g., Acharya et al., 2011; Bhatt & Joshi, 2011; Price et al., 2014; Elsen et al., 2017; Srinivasan et al., 2018; Schumm et al., 2020). While these are relatively recent, unlike data from the Andes and from Papua New Guinea, in time, these data can be used as a baseline to compare later-day elevational ranges.

A further promising area for research is to tease apart differences between abiotic factors such as temperature, biotic interactions such as competition and resource use and factors such as vegetation structure in influencing the elevational ranges of species (Elsen et al. 2017, Burner et al., 2020). These factors will likely have different roles in influencing the ranges of different species and populations (e.g., frugivores might be more expected to be constrained by the distribution of fruiting plants), thereby also influencing how species traits such as body size and diet interact with climate change to reset elevational ranges. The rate of shift of whole habitats and how these lag behind climate may also impact the continued survival of species; for example, while Bornean moths shift upward fairly concurrently with warming (Wu et al., 2019), plants in the Alps tend to lag far behind (Rumpf et al., 2018), though this could be a consequence of the long generation time of trees coupled with the inherent thermal tolerances of these plant species. This creates the problem of the movements of mobile taxa tied to a specialized habitat being constrained by slow upward shifts of vegetation compared to climate. Local topographies (Gaüzère et al., 2017) and increasing seasonality with climate change and consequent shifts in hygric niches (Boyle et al., 2020) could also impact the populations independent of upslope shifts; for instance, local microclimatic refugia against more pronounced seasonality with further climate warming might in fact allow species to survive for longer durations at a certain elevation without upslope shifts. Hence, building a complete understanding of species’ responses to climate change is important to predict which kinds of species are most vulnerable to continued warming in tropical mountains, and to identify potential steps to minimise the impacts of climate change on such species. In the eastern Himalayas, it is likely that birds will require contiguous expanses of old-growth forest across elevational gradients to enable upslope range shifts and maintain resilience to climate change (Srinivasan & Wilcove 2020).

One future direction of study would be to compare these rates of upslope shift for tropical montane biotas across the planet, in order to assess which regions and taxonomic groups are most at risk to possible extirpations, or even extinction in the case of range-restricted endemic species (Carroll et al., 2015). This preliminary prediction technique is a tractable framework for making inferences in other tropical biotas, prerequisite to the presence of a significant number of citizen science datasets for the region, and information on the elevational bounds for at least some of the species recorded there.

## ACKNOWLEDGEMENTS

We thank all the birdwatchers who uploaded their records from Eaglenest Wildlife Sanctuary and Singchung Bugun Village Community Reserve to eBird, and for eBird for making such a platform available.

## Ethics Statement

No animals were handled as part of this research.

## Author Contributions

K.S.G. and U.S. conceptualized the study, K.S.G and U.S. designed the methods of analysis and analyzed the data, K.S.G. wrote the first draft, U.S. provided revisions and additions to the manuscript.

## Data Depository

The data as well as the code used in this analysis will be added on DataDryad after acceptance in a journal and prior to publication.

## LITERATURE CITED

Acharya, B. K., Sanders, N. J., Vijayan, L., & Chettri, B. (2011). Elevational gradients in bird diversity in the Eastern Himalaya: an evaluation of distribution patterns and their underlying mechanisms. PloS One, 6(12), e29097.

Angert, A. L., Crozier, L. G., Rissler, L. J., Gilman, S. E., Tewksbury, J. J., & Chunco, A. J. (2011). Do species’ traits predict recent shifts at expanding range edges? Ecology Letters, 14(7), 677–689.

Billerman, S. M., Keeney, B. K., Rodewald, P. G., and Schulenberg, T. S. (Editors) (2020). Birds of the World. Cornell Laboratory of Ornithology, Ithaca, NY, USA. https://birdsoftheworld.org/bow/home

Bhatt, D., & Joshi, K. K. (2011). Bird assemblages in natural and urbanized habitats along elevational gradient in Nainital district (Western Himalaya) of Uttarakhand state, India. Current Zoology, 57(3), 318–329.

Boyle, W. A., Shogren, E. H., & Brawn, J. D. (2020). Hygric niches for tropical endotherms. Trends in Ecology & Evolution.

Brook, B. W., Sodhi, N. S., & Bradshaw, C. J. (2008). Synergies among extinction drivers under global change. Trends in Ecology & Evolution, 23(8), 453–460.

Burner, R. C., Styring, A. R., Rahman, M. A., & Sheldon, F. H. (2019). Occupancy patterns and upper range limits of lowland Bornean birds along an elevational gradient. Journal of Biogeography, 46(11), 2583–2596.

Burner, R. C., Boyce, A. J., Bernasconi, D., Styring, A. R., Shakya, S. B., Boer, C., … & Sheldon, F. H. (2020). Biotic interactions help explain variation in elevational range limits of birds among Bornean mountains. Journal of Biogeography, 47(3), 760–771.

Carroll, C., Lawler, J. J., Roberts, D. R., & Hamann, A. (2015). Biotic and climatic velocity identify contrasting areas of vulnerability to climate change. PloS One, 10(10), e0140486.

Ceballos, G., & Ehrlich, P. R. (2018). The misunderstood sixth mass extinction. Science, 360(6393), 1080.2–1081

Chen, I.-C., Shiu, H.-J., Benedick, S., Holloway, J. D., Chey, V. K., Barlow, H. S., Hill, J. K., & Thomas, C. D. (2009). Elevation increases in moth assemblages over 42 years on a tropical mountain. Proceedings of the National Academy of Sciences, 106(5), 1479–1483.

Chen, I.-C., Hill, J.K., Ohlemüller, R., Roy, D.B., Thomas, C.D., (2011). Rapid range shifts of species associated with high levels of climate warming. Science 333, 1024–1026

eBird. 2020. eBird: An online database of bird distribution and abundance [web application]. eBird, Cornell Lab of Ornithology, Ithaca, New York. Available: http://www.ebird.org. (June 22, 2020).

Elsen, P. R., & Tingley, M. W. (2015). Global mountain topography and the fate of montane species under climate change. Nature Climate Change, 5(8), 772–776.

Elsen, P. R., Tingley, M. W., Kalyanaraman, R., Ramesh, K., & Wilcove, D. S. (2017). The role of competition, ecotones, and temperature in the elevational distribution of Himalayan birds. Ecology, 98(2), 337–348.

Feeley, K. J., Silman, M. R., Bush, M. B., Farfan, W., Cabrera, K. G., Malhi, Y., … & Saatchi, S. (2011). Upslope migration of Andean trees. Journal of Biogeography, 38(4), 783–791.

Freeman, B. G., & Freeman, A. M. C. (2014). Rapid upslope shifts in New Guinean birds illustrate strong distributional responses of tropical montane species to global warming. Proceedings of the National Academy of Sciences, 111(12), 4490–4494.

Freeman, B. G., Scholer, M. N., Ruiz-Gutierrez, V., & Fitzpatrick, J. W. (2018). Climate change causes upslope shifts and mountaintop extirpations in a tropical bird community. Proceedings of the National Academy of Sciences, 115(47), 11982–11987.

Freeman, B. G., Tobias, J. A., & Schluter, D. (2019). Behavior influences range limits and patterns of coexistence across an elevational gradient in tropical birds. Ecography, 42(11), 1832–1840.

Freeman, B. G., Song, Y., Feeley, K., & Zhu, K. (2020). Montane species and communities track recent warming more closely in the tropics. bioRxiv: doi.org/10.1101/2020.05.18.102848

Gaston, K. J. (2009). Geographic range limits: achieving synthesis. Proceedings of the Royal Society B: Biological Sciences, 276(1661), 1395–1406.

Gaüzère, P., Princé, K., & Devictor, V. (2017). Where do they go? The effects of topography and habitat diversity on reducing climatic debt in birds. Global Change Biology, 23(6), 2218–2229.

Grenyer, R., Orme, C. D. L., Jackson, S. F., Thomas, G. H., Davies, R. G., Davies, T. J., … & Ding, T. S. (2006). Global distribution and conservation of rare and threatened vertebrates. Nature, 444(7115), 93–96.

Intergovernmental Panel on Climate Change (IPCC) (2007). Fourth Assessment Report: Climate Change 2007, The Physical Science Basis. Cambridge University Press, Cambridge.

Janzen, D. H. (1967). Why mountain passes are higher in the tropics. The American Naturalist, 101(919), 233–249.

La Sorte, F. A., & Jetz, W. (2010). Projected range contractions of montane biodiversity under global warming. Proceedings of the Royal Society B: Biological Sciences, 277(1699), 3401–3410.

Lenoir, J., & Svenning, J. C. (2015). Climate-related range shifts–a global multidimensional synthesis and new research directions. Ecography, 38(1), 15–28.

Lenoir, J., Bertrand, R., Comte, L., Bourgeaud, L., Hattab, T., Murienne, J., & Grenouillet, G. (2020). Species better track climate warming in the oceans than on land. Nature Ecology & Evolution, 1–16.

Marris, E. (2007). The escalator effect. Nature Reports: Climate Change, 1, 94–96.

Moritz, C., Patton, J. L., Conroy, C. J., Parra, J. L., White, G. C., & Beissinger, S. R. (2008). Impact of a century of climate change on small-mammal communities in Yosemite National Park, USA. Science, 322(5899), 261–264.

Morueta-Holme, N., Engemann, K., Sandoval-Acuña, P., Jonas, J. D., Segnitz, R. M., & Svenning, J. C. (2015). Strong upslope shifts in Chimborazo’s vegetation over two centuries since Humboldt. Proceedings of the National Academy of Sciences, 112(41), 12741–12745.

Myers, N., Mittermeier, R. A., Mittermeier, C. G., Da Fonseca, G. A., & Kent, J. (2000). Biodiversity hotspots for conservation priorities. Nature, 403(6772), 853–858.

Parmesan, C., & Yohe, G. (2003). A globally coherent fingerprint of climate change impacts across natural systems. Nature, 421(6918), 37–42.

Perez T. M., Stroud, J. T. & Feeley, K. J. (2016). Thermal trouble in the tropics. Science 351, 1392–1393.

Pimm, S. L. (2008). Biodiversity: climate change or habitat loss—which will kill more species?. Current Biology, 18(3), R117–R119.

Price, T. D., Hooper, D. M., Buchanan, C. D., Johansson, U. S., Tietze, D. T., Alström, P., … & Martens, J. (2014). Niche filling slows the diversification of Himalayan songbirds. Nature, 509(7499), 222–225.

R Core Team (2020). R: A language and environment for statistical computing. R Foundation for Statistical Computing, Vienna, Austria. URL: https://www.R-project.org/.

Raxworthy, C. J., Pearson, R. G., Rabibisoa, N., Rakotondrazafy, A. M., Ramanamanjato, J. B., Raselimanana, A. P., … & Stone, D. A. (2008). Extinction vulnerability of tropical montane endemism from warming and upslope displacement: a preliminary appraisal for the highest massif in Madagascar. Global Change Biology, 14(8), 1703–1720.

Rowe, K. C., Rowe, K. M., Tingley, M. W., Koo, M. S., Patton, J. L., Conroy, C. J., … & Moritz, C. (2015). Spatially heterogeneous impact of climate change on small mammals of montane California. Proceedings of the Royal Society B: Biological Sciences, 282(1799), 20141857.

Rumpf, S. B., Hülber, K., Klonner, G., Moser, D., Schütz, M., Wessely, J., … & Dullinger, S. (2018). Range dynamics of mountain plants decrease with elevation. Proceedings of the National Academy of Sciences, 115(8), 1848–1853.

Salick, J., Fang, Z., & Hart, R. (2019). Rapid changes in eastern Himalayan alpine flora with climate change. American Journal of Botany, 106(4), 520–530.

Schumm, M., White, A. E., Supriya, K., & Price, T. D. (2020). Ecological limits as the driver of bird species richness patterns along the east Himalayan elevational gradient. The American Naturalist, 195(5), 802–817.

Şekercioğlu, Ç. H., Schneider, S. H., Fay, J. P., & Loarie, S. R. (2008). Climate change, elevational range shifts, and bird extinctions. Conservation Biology, 22(1), 140–150.

Şekercioğlu, Ç. H., Primack, R. B., & Wormworth, J. (2012). The effects of climate change on tropical birds. Biological Conservation, 148(1), 1–18.

Shrestha, U. B., Gautam, S., & Bawa, K. S. (2012). Widespread climate change in the Himalayas and associated changes in local ecosystems. PloS One, 7(5), e36741.

Spierenburg, P. (2005). Birds in Bhutan: Status and Distribution. Oriental Bird Club, 1–383.

Srinivasan, U., Elsen, P. R., Tingley, M. W., & Wilcove, D. S. (2018). Temperature and competition interact to structure Himalayan bird communities. Proceedings of the Royal Society B: Biological Sciences, 285, 20172593.

Srinivasan, U., & Wilcove, D.S. (2020). Interactive impacts of climate change and land-use change on the demography of montane birds. In press, Ecology.

Thomas, C. D., Cameron, A., Green, R. E., Bakkenes, M., Beaumont, L. J., Collingham, Y. C., … & Hughes, L. (2004). Extinction risk from climate change. Nature, 427(6970), 145–148.

Wormworth, J., & Şekercioğlu, Ç. H. (2011). Winged sentinels: birds and climate change. Cambridge University Press.

Wu, C. H., Holloway, J. D., Hill, J. K., Thomas, C. D., Chen, I. C., & Ho, C. K. (2019). Reduced body sizes in climate-impacted Borneo moth assemblages are primarily explained by range shifts. Nature Communications, 10.

